# Is faster always better? The walking speed-dependency of gait variability in bilateral vestibulopathy

**DOI:** 10.1101/413955

**Authors:** Christopher McCrum, Florence Lucieer, Raymond van de Berg, Paul Willems, Angélica Pérez Fornos, Nils Guinand, Kiros Karamanidis, Herman Kingma, Kenneth Meijer

## Abstract

Study of balance and gait deficits associated with vestibulopathy is important for improving clinical care and is critical to our understanding of the vestibular contributions to gait and balance control. Previous studies report a speed-dependency of the vestibular contributions to gait, so we examined the walking speed effects on gait variability in healthy young and older adults and in adults with bilateral vestibulopathy (BVP). Forty-four people with BVP, 12 healthy young adults and 12 healthy older adults completed walking trials at 0.4m/s to 1.6m/s in 0.2m/s intervals on a dual belt, instrumented treadmill. Using a motion capture system and kinematic data, the means and coefficients of variation for step length, time, width and double support time were calculated. The BVP group also completed a video head impulse test and examinations of ocular and cervical vestibular evoked myogenic potentials and dynamic visual acuity. Walking speed significantly affected all assessed gait parameters. Step length variability at slower speeds and step width variability at faster speeds were the most distinguishing parameters between the healthy participants and people with BVP, and within people with BVP with different locomotor capacities. We observed for step width variability, specifically, an apparent persistent importance of vestibular function at increasing speeds. Gait variability was not associated with the clinical vestibular tests. Our results indicate that gait variability at multiple walking speeds has potential as an assessment tool for vestibular interventions.

**New & Noteworthy:** Walking speed significantly but differentially affects gait variability in healthy adults and in adults with bilateral vestibulopathy. Gait variability at different speeds distinguishes between participants with and without bilateral vestibulopathy, but also between more and less able walkers with bilateral vestibulopathy. Specifically, for step width variability, an apparent persistent importance of vestibular function at increasing walking speeds was observed. Gait variability was generally not correlated with clinical tests of vestibular function.

## Introduction

Ever since a chance observation of a dog with acute unilateral vestibulopathy who demonstrated less imbalance during running than during walking (Brandt et al. 1999), the interactions of gait velocity, imbalance and vestibular symptoms in people with vestibulopathy have become a topic of great interest. Inspired by the observation in the dog, Brandt et al. (1999) demonstrated with a simple setup that humans with acute unilateral vestibulopathy could run with less deviation to the affected side than while walking. Since then, three studies have reported reductions in temporal gait variability and reductions in stride length variability in bilateral vestibulopathy (BVP) during faster, compared to slower walking (Schniepp et al. 2017; Schniepp et al. 2012; Wuehr et al. 2016). BVP, a severe bilateral reduction of vestibular function that results in severe balance deficits and an increased risk of falls (Guinand et al. 2012a; Horak et al. 2016; Lucieer et al. 2016; Schlick et al. 2016; Sprenger et al. 2017; van de Berg et al. 2015), was recently defined by the Bárány Society (Strupp et al. 2017) and represents one of the most debilitating vestibular disorders. Interestingly, the same studies revealed that patients with BVP do not self-select walking speeds that minimize temporal or spatial gait variability (Schniepp et al. 2017; Schniepp et al. 2012; Wuehr et al. 2016), which may suggest that these are not the only source of instability or inefficiency with which people with BVP must cope. The study of the severe balance and gait deficits in people with BVP is both important for improving clinical care and for objective quantification of the effects of novel interventions, such as vestibular implants (Guyot et al. 2016; Lewis 2016). Furthermore, it is fundamental to our understanding of the vestibular contributions to gait and balance control.

The sensory contributions to gait appear to depend on walking speed, which may partly explain the above described findings and will affect walking speed selection in people with vestibulopathy. Visual perturbations such as distorting prisms or closure of the eyes have less impact on most gait variability parameters the faster one walks (Jahn et al. 2001; Wuehr et al. 2013) with the exception of step width variability, which appears to increase with visual perturbation at faster walking speeds (Wuehr et al. 2013). Similarly, vestibular perturbations via galvanic vestibular stimulation have less impact on gait direction and variability at higher speeds (Fitzpatrick et al. 1999; Jahn et al. 2000). It has also been reported that the vestibular influence on lower limb muscles (determined by examining vestibulo-muscular coupling via lower limb muscle electromyography during vestibular stimulation) is selectively suppressed with increased cadence and speed during walking (Dakin et al. 2013; Forbes et al. 2017), purported to be related to a shift in the control mechanisms of mediolateral stability with increasing walking speeds from active stabilization at the lower limb joints during the stance phase to foot placement (Bauby and Kuo 2000; Dakin et al. 2013). Despite selective suppression of the vestibular influence on some lower limb muscles at faster walking speeds, significant increases in frontal spatial variability with increasing walking speeds have been reported in BVP (Wuehr et al. 2016), suggesting that vestibular information remains important for mediolateral stability during gait at faster speeds.

In order to further investigate the walking speed dependency of gait variability in vestibulopathy, this study analyzed the gait of people with BVP and of healthy control participants. We aimed to determine the effects of systematic increases in walking speed on spatiotemporal gait parameters and their variability in these participant groups. Secondly, we aimed to assess if these parameters would differentiate between healthy participants, and participants with BVP who could and could not complete all of the planned walking speed trials (used here as a simple proxy of locomotor capacity; see Methods). We hypothesized that, for all participants, step and double support time and step length variability would systematically reduce with increases in walking speed, whereas step width variability would systematically increase, in agreement with previous work (Schniepp et al. 2017; Schniepp et al. 2012; Wuehr et al. 2016). We further postulated that step and double support time and step length variability at slower walking speeds would be most distinguishing between the healthy control participants and patients with BVP, and also between the patients with BVP that could complete the measurement protocol, and the patients with BVP that could only partially complete the measurement protocol, whereas step width variability would be most distinguishing at faster walking speeds. Additionally, we aimed to conduct an explorative analysis in the patient groups by examining correlations between the outcomes of the most distinguishing gait parameters identified and clinical vestibular tests (video head impulse test [vHIT], ocular and cervical vestibular evoked myogenic potentials [oVEMP and cVEMP] and dynamic visual acuity [DVA]) that are indicative of vestibular functional integrity.

## Materials and Methods

### Participants

Forty four people with BVP participated in this study (22 males, 22 females; age: 57.6±11.5 years, age range: 21 to 74; height: 174.5±9.7cm; weight: 80.4±17kg). Inclusion criteria were a prior diagnosis of bilateral vestibular hypofunction at the Maastricht University Medical Centre+ (imbalance and/or oscillopsia during locomotion and summated slow phase mean peak velocity of the nystagmus of less than 20°/s during bithermal caloric tests) and the self-reported ability to walk independently without assistance. Please note that this study began prior to the publication of the Bárány Society guidelines (Strupp et al. 2017), which are slightly different. Potential participants were not included if they were unable or unwilling to stop taking anxiety or depression medication for the week before the measurements. In addition, two healthy control groups comprised of 12 healthy younger adults (Young; 5 males, 7 females; 25.1±2.8 years; 174.9±7.3cm; 72.6±13.5kg) and 12 healthy older adults (Older; 8 males, 4 females; 71.5±4.8 years; 171.5±9.1cm; 79.5±11.8kg) with no history of balance or gait difficulties and no history of dizziness participated in this study. These specific groups were included to account for the age range in the BVP group and to provide an estimation of the effect of ageing alone on the outcome parameters. The study was explained before obtaining written informed consent, was conducted in accordance with the Declaration of Helsinki and was approved by the Maastricht University Medical Centre medical ethics committee (gait measurements: NL58205.068.16; vestibular tests: NL52768.068.15).

### Gait Analysis Setup, Data Processing and Procedure

The gait measurements were conducted using the Computer Assisted Rehabilitation Environment Extended (CAREN; Motekforce Link, Amsterdam, The Netherlands), which includes a dual-belt force plate-instrumented treadmill (Motekforce Link, Amsterdam, The Netherlands; 1000Hz), a 12 camera motion capture system (100Hz; Vicon Motion Systems, Oxford, UK) and a virtual environment (city-style street with passing objects and structures) projected onto a 180 degrees curved screen (note that optic flow was turned off for the participants with BVP to prevent dizziness and nausea). For all measurement sessions, a safety harness connected to an overhead frame was used. At the request of some of the participants with BVP, a handrail was also positioned on the treadmill, the use of which was monitored and recorded. Six retroreflective markers were attached to anatomical landmarks (C7, sacrum, left and right trochanter and left and right hallux) and were tracked by the motion capture system. Marker tracks were filtered using a low pass second order Butterworth filter (zero-phase) with a 12Hz cut-off frequency. Foot touchdown was determined using combined force plate (50N threshold) and foot marker data (Zeni et al. 2008). This combined method was used to be able to accurately account for foot touchdowns and toe-offs occurring in the center of the treadmill triggering both force plates simultaneously. For these steps, the foot marker method was used and then corrected based on the average discrepancy between the force plate method and the marker method timing for all steps that contacted only one force plate. The spatiotemporal gait parameters of interest were step length (anteroposterior distance between the hallux markers at foot touchdown), step time (time from touchdown of one foot to touchdown of the next foot), step width (mediolateral distance between the hallux markers at foot touchdown) and double support time (time spent with both feet on the ground). Means, standard deviations and coefficients of variation (CV) were determined for each speed for each participant.

Each session began with walking familiarization trials at 0.4m/s up to 1.6m/s in 0.2m/s intervals. At least 60s were used for each speed, and further time was provided to familiarize to each speed if deemed necessary by either the participant, the CAREN operator or the research clinician. At the end of each speed trial, the decision to continue to the next (faster) speed was made in a similar manner. If the participant was not comfortable progressing to the next speed or if the CAREN operator or research clinician did not think it was safe or feasible to progress, then the participant continued at the current speed instead. Participants were then given sufficient rest before continuing with the measurements. Single two-to-three-minute-long measurements (to ensure a minimum of 60 strides per speed) were then conducted at each prescribed speed that was completed during familiarization. Multiple set walking speeds were used as opposed to the majority of previous studies which have used either percentages of preferred walking speeds or self-perceived slow, normal and fast walking speeds, in order to have more control over the walking speed condition.

### Clinical Vestibular Function Tests Setup and Procedures

Following a sufficient rest period that was determined on an individual basis, the BVP group proceeded with the clinical vestibular testing battery. Between each test, sufficient rest was provided based on feedback from the patient and the judgement of the clinical researcher. The vHIT was performed with the EyeSeeCam system (EyeSeeCam VOG; Munich, Germany) and the ICS Impulse system (GN Otometrics A/S, Denmark). Both systems measured the movement of the right eye. The distance of the back of the static chair was 2 meters to the point of fixation. The point of fixation consisted of a green dot on the wall, produced by a laser on a tripod. If necessary, adhesive plasters were used to lift the upper eyelid a little to secure the visibility of the pupil for the camera in all directions. Goggle movement was minimized by adjusting the strap of the goggles to every subject. The vHIT system was calibrated according to the protocol of the system. After calibration, the subject was instructed to not touch their head including the goggles. The examiner stood behind the participant with two hands firmly on top of the participant’s head without touching the strap of the goggles. The examiner then applied head impulses in six different movements to test each canal (McGarvie et al. 2015). The horizontal head impulses comprised a peak velocity of > 150°/s and the vertical head impulses a peak head velocity of > 100°/s. The amplitude of the movements was 10-20°. Only outward impulses were used (van Dooren et al. 2018). The vHIT was defined as abnormal if the VOR-gain was below 0,7 and/or if covert saccades were observed in 50% or more of the traces (McGarvie et al. 2015; Yip et al. 2016).

DVA was assessed on a regular treadmill (1210 model, SportsArt, Inc., Tainan, Taiwan, China.) with the participant positioned 2.8 meters from a computer screen. Firstly, the static visual acuity was determined during stance, followed by the assessment of the DVA during walking at 2, 4 and 6 km/h. One letter at a time was randomly displayed on the screen from a chart of Sloan letters (CDHKNORSVZ; Sloan 1959). Starting at a logMAR (log of the Minimum Angle of Resolution; (Bailey and Lovie 1976)) of 1.0, five random letters were shown at each logMAR (decreasing in steps of 0.1 logMAR). When four out of five letters were correctly identified, the corresponding logMAR was considered achieved. The outcome of the DVA was the difference between the static logMAR and the logMAR for each of the three walking speeds. The result was omitted if the subject needed a handrail to walk at that speed or if it wasn’t possible to walk at that speed at all (Guinand et al. 2012b).

cVEMP and oVEMP were assessed with the Neuro-Audio system (v2010, Neurosoft, Ivanovo, Russia). A monaural stimulation with in-ear earphones was used with air conduction tone bursts at 500Hz and a stimulation rate of 13Hz using a blackman window function with a two-cycle rise/fall and no plateau phase. Tone bursts of maximum 130dB sound pressure level (SPL) were used. A stepwise approach was used to determine the threshold with a precision of 5dB SPL (van Tilburg et al. 2016). Positive (P1) and negative (N1) peaks in the recorded biphasic waveform were marked for both cVEMPs and oVEMPs. The thresholds were determined as the lowest stimulus intensities to elicit recognizable peaks. If it wasn’t possible to find a VEMP response, it was defined as a threshold of >130dB SPL. For the cVEMP, the participant was positioned lying down with the back positioned at a 30° angle above the horizontal plane and was asked to turn their head towards the non-measured side and lift their head during the measurement. The cVEMP was recorded at the ipsilateral sternocleidomastoid muscle. Two electrodes were placed on the sternocleidomastoid muscles, the reference electrode on the sternum, and the earth electrode on the forehead. Electrode impedances of 5 kΩ or lower were accepted and otherwise the electrode was replaced. To ensure correct muscle contraction, a feedback system using a screen was provided. An average of 200 EMG traces with a minimum mean rectified voltage (MRV) of 65µV and a maximum MRV of 205µV was accepted (Brantberg and Lofqvist 2007; Fujimoto et al. 2009). The oVEMP was recorded at the contralateral inferior oblique muscle. Five electrodes were used: the recording electrodes beneath the eyelid, just lateral of the pupil when gazing forward and centrally, the reference electrodes beneath the recording electrode and the earth electrode on the forehead. The participant was asked to keep their gaze at a focus point placed at a 30 degrees angle behind the head. An average of at least 300 EMG traces was accepted (Govender et al. 2011; Piker et al. 2013; Valko et al. 2016).

### Statistics

From the 44 participants with BVP that started the study, 38 participants were able to complete at least the three slowest walking speeds without assistance (group hereafter referred to as BVP) and these participants’ data were taken for the comparison with the healthy groups. For the within BVP comparisons, three groups were formed. One group was able to complete all of the gait measurements without assistance (BVP All Gait; n=26), the second was only able to complete some of the speeds without assistance (BVP Part Gait; n=12; all of this group were able to complete the measurements at least up to 0.8m/s) and the final group (BVP No Gait; n=6) did not start the recorded gait trials (see “Results” for details on this group).

To investigate the effects of walking speed on gait and this effect’s potential interaction with vestibular function, mixed-effects models using the restricted maximum likelihood method with the fixed effects walking speed, participant group, and speed by group interaction were conducted for the means and CVs of step time and length, step width and double support time. To further investigate the potential of gait variability to distinguish between BVP groups, mixed-effects models as described above were applied with groups BVP All Gait and BVP Part Gait to the CV of all four gait parameters across all speeds that included data points from each group. Bonferroni post hoc comparisons were performed to assess the group differences within speeds for each of the gait parameters.

The vHIT testing revealed abnormal canal function in all or most directions for almost all of the participants with BVP (i.e. exceptions were two participants with BVP who had only one abnormal result out of six). As almost all outcomes were abnormal and there was no possibility to distinguish between groups, analysis of the vHIT results in relation to gait was not taken further. For all completed DVA trials with a logMAR change value during the three walking speeds compared to standing and when oVEMP or cVEMP thresholds were detected, these values were grouped and Pearson correlations with the gait parameters that showed highest variability and/or distinguished between BVP groups were conducted (see Results). Age, height, weight and body mass index (BMI) were compared across the participant groups BVP, Young and Older, and within the three BVP groups (BVP All Gait, BVP Part Gait, BVP No Gait) using one way ANOVAs with Bonferroni corrections for multiple comparisons.

## Results

Twenty six participants with BVP were able to complete all of the gait measurements without assistance (BVP All Gait). Twelve participants with BVP were only able to complete some of the speeds (BVP Part Gait), of which one participant stopped after 0.8m/s, one after 1.0m/s, four after 1.2m/s and six after 1.4m/s. Six participants with BVP were assigned to the BVP No Gait group for the following reasons: one participant became dizzy and nauseated during familiarization and could not continue; three participants were not able to walk during familiarization without handrail support; two participants found treadmill walking too challenging and could not continue. The demographic data of these three groups, as well as the healthy control group can be found in Table 1. The one-way ANOVAs revealed a significant group effect (BVP, Young, Older) for age (F (2,59) = 88), P<0.0001), with age significantly differing between each of the groups (P<0.0001). Height, weight and BMI did not significantly differ across these groups. No significant differences in demographics were found with the three BVP groups.

**Table 1.** Participant Group Characteristics

|  | **n** | **Age (y)** | **Height (cm)** | **Weight (kg)** | **Body Mass Index** |
| --- | --- | --- | --- | --- | --- |
| **Young** | 12 (7 female) | 25.1±2.8* | 174.9±7.3 | 72.6±13.5 | 23.6±2.8 |
| **Older** | 12 (4 female) | 71.5±4.8* | 171.5±9.1 | 79.5±11.8 | 26.9±2.2 |
| **BVP** | 38 (20 female) | 56.1±11* | 174.6±10.1 | 80.2±17.6 | 26.1±4.2 |
| ***BVP All Gait*** | 26 (10 female) | 55.1±11.4 | 176.8±9.9 | 80.3±17.8 | 25.4±3.8 |
| ***BVP Part Gait*** | 12 (10 female) | 59.2±9 | 169.7±9 | 79.9±18 | 27.6±4.7 |
| ***BVP No Gait*** | 6 (2 female) | 65.3±13.6 | 174±6.9 | 82.4±13.4 | 27.2±3.8 |

Values are means ± SD. \*: Significantly different from each other ( $P < 0.0001$ ).

The mixed-effects models with walking speed (0.4 to 1.6m/s) and group (BVP, Young, Older) as factors revealed significant walking speed effects for the means and CV of step time and length, step width and double support time (*P*≤0.0003), significant group effects for all parameters except step width means (*P*≤0.0151) and significant walking speed by group interactions for the means of step time, double support time and step width (*P*≤0.0053) and the CV of step width (*P*<0.0001). The mixed-effects model results and summary of the between group Bonferroni comparisons are displayed in Fig. 1 (means) and Fig. 2 (CVs), and the full Bonferroni comparison results are available in Supplementary Tables 1 and 2.

**Fig. 1.**
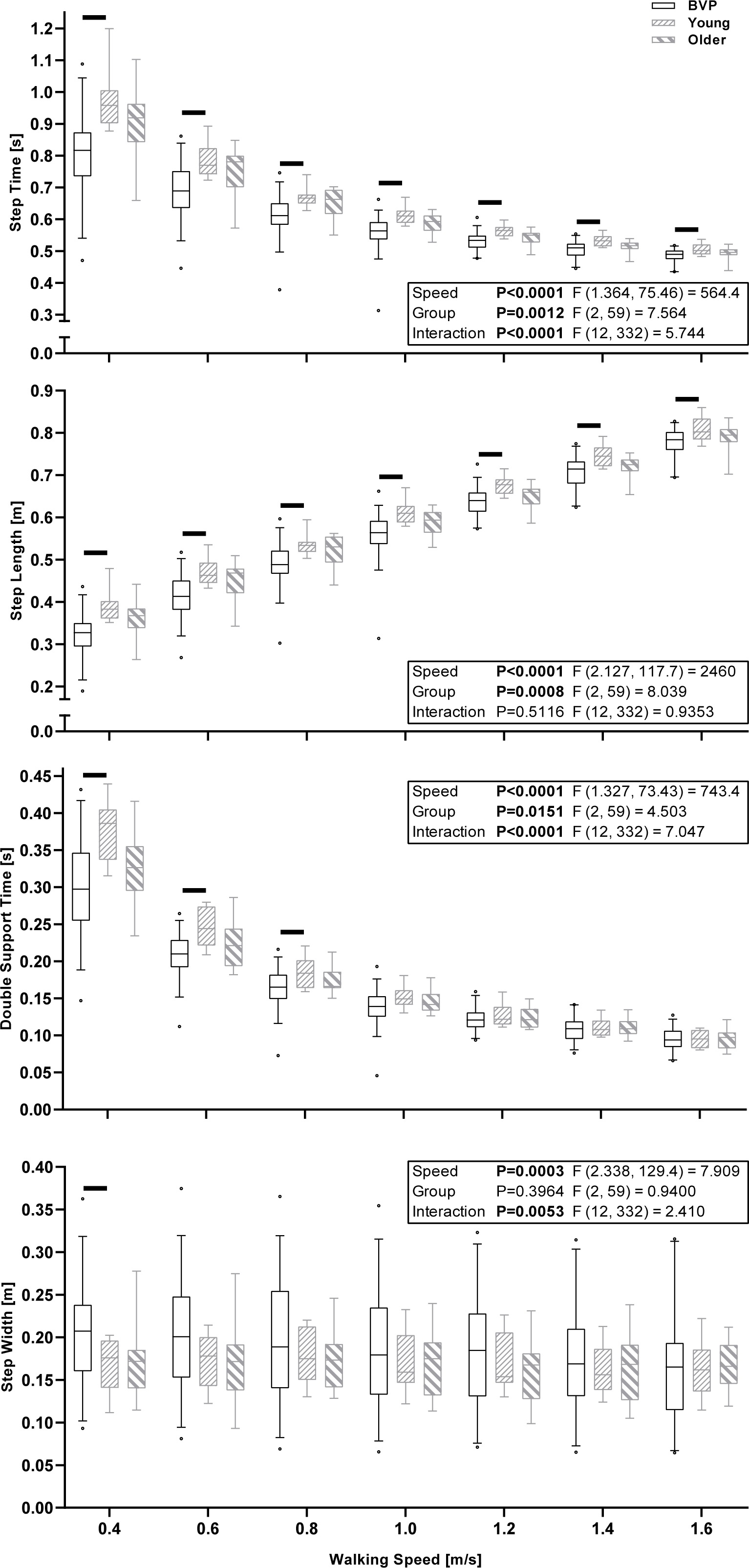
Boxplots of the median, interquartile range and 5^th^ and 95^th^ percentile of the means of step time, step length, double support time and step width across all conducted walking speeds in BVP, Young and Older participant groups. The black horizontal lines indicate significant between group differences for the indicated speed (P<0.05, Bonferroni adjusted).

**Fig. 2.**
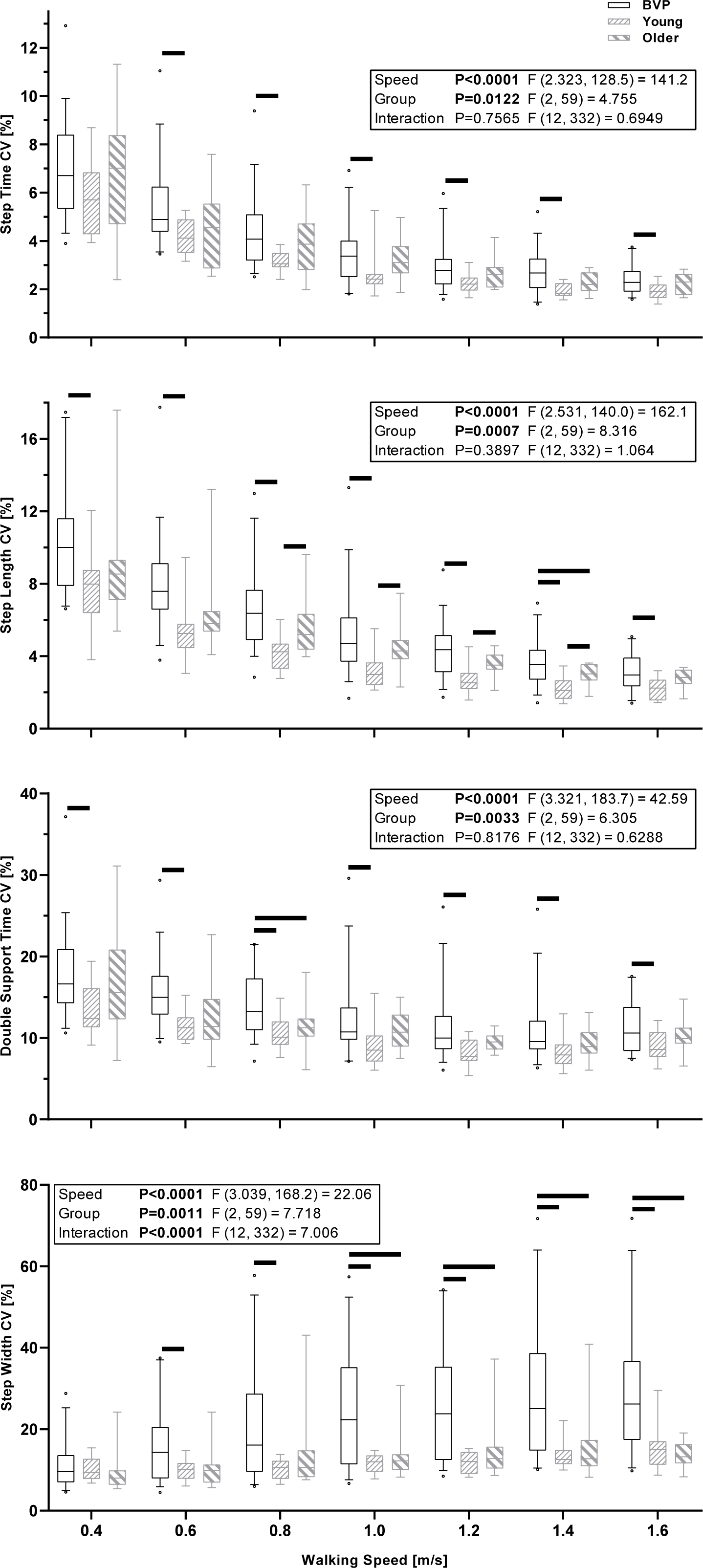
Boxplots of the median, interquartile range and 5^th^ and 95^th^ percentile of the coefficients of variation (CV) of step time, step length, double support time and step width across all conducted walking speeds in BVP, Young and Older participant groups. The black horizontal lines indicate significant between group differences for the indicated speed (P<0.05, Bonferroni adjusted).

The mixed-effects models with walking speed (0.4 to 1.4m/s) and group (BVP All Gait and BVP Part Gait) as factors revealed significant effects of walking speed for the CV of all parameters (*P*<0.0001). Significant group effects were found for the CV of step time, step length and double support time (*P*≤0.0162) and a significant walking speed by group interaction was found for the CV of double support time (*P*=0.0172). The mixed-effects model results and summary of the between group Bonferroni comparisons are displayed in Fig. 3 and the full Bonferroni comparison results are available in Supplementary Table 3.

**Fig. 3.**
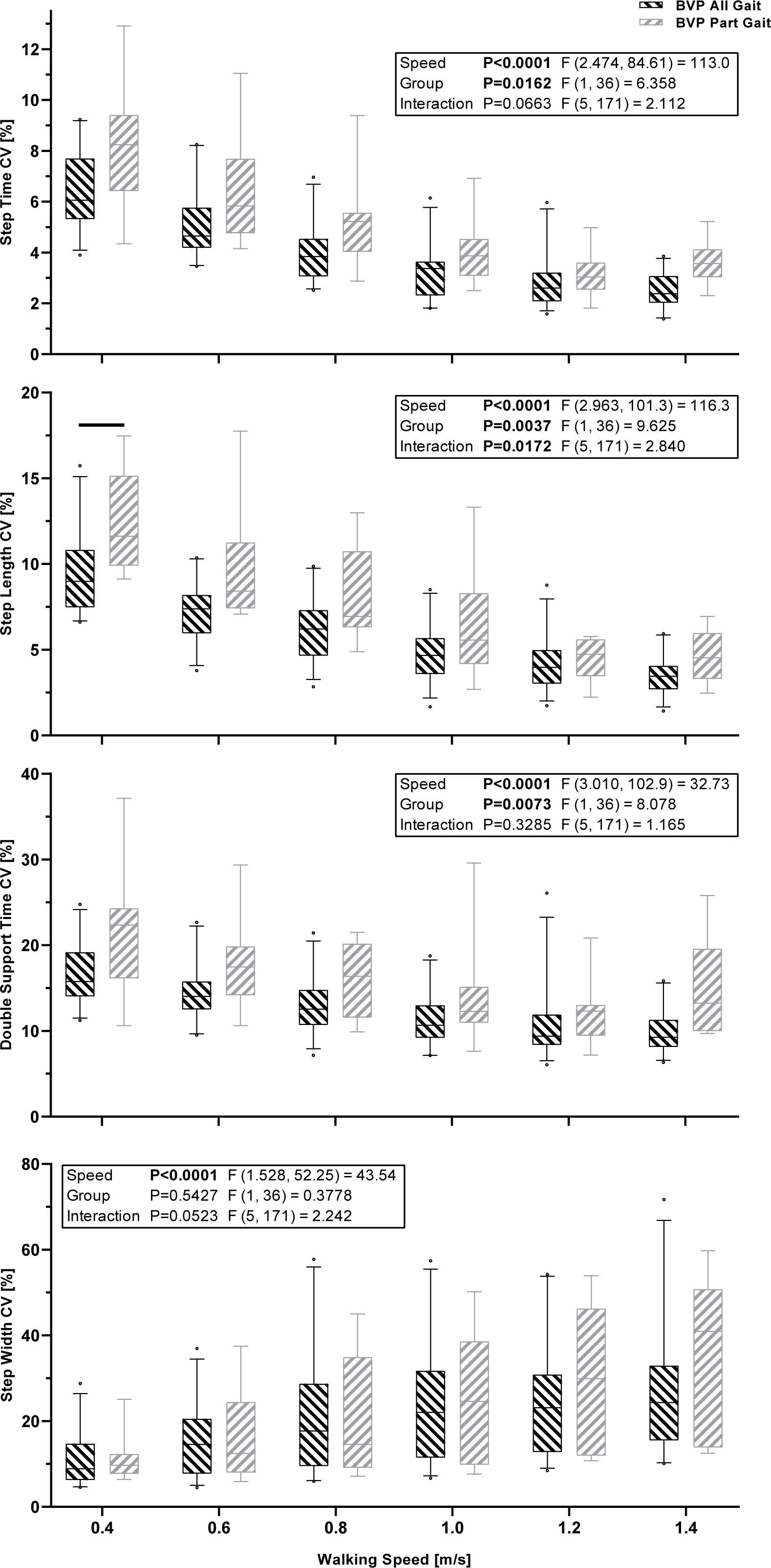
Boxplots of the median, interquartile range and 5^th^ and 95^th^ percentile of the coefficients of variation (CV) of step time, step length, double support time and step width across all walking speeds with data from participant groups BVP All Gait and BVP Part Gait. The black horizontal lines indicate significant between group differences for the indicated speed (P<0.05, Bonferroni adjusted).

When cVEMP and oVEMP thresholds were detected, and when a speed of the DVA was completed, these values were taken and Pearson correlations were conducted with the CVs of step time, step length and double support time at 0.4m/s and the CV of step width at 1.6m/s, being the speeds with the highest variability in those parameters from the previous analysis. These results can be seen in Table 2.

**Table 2:**
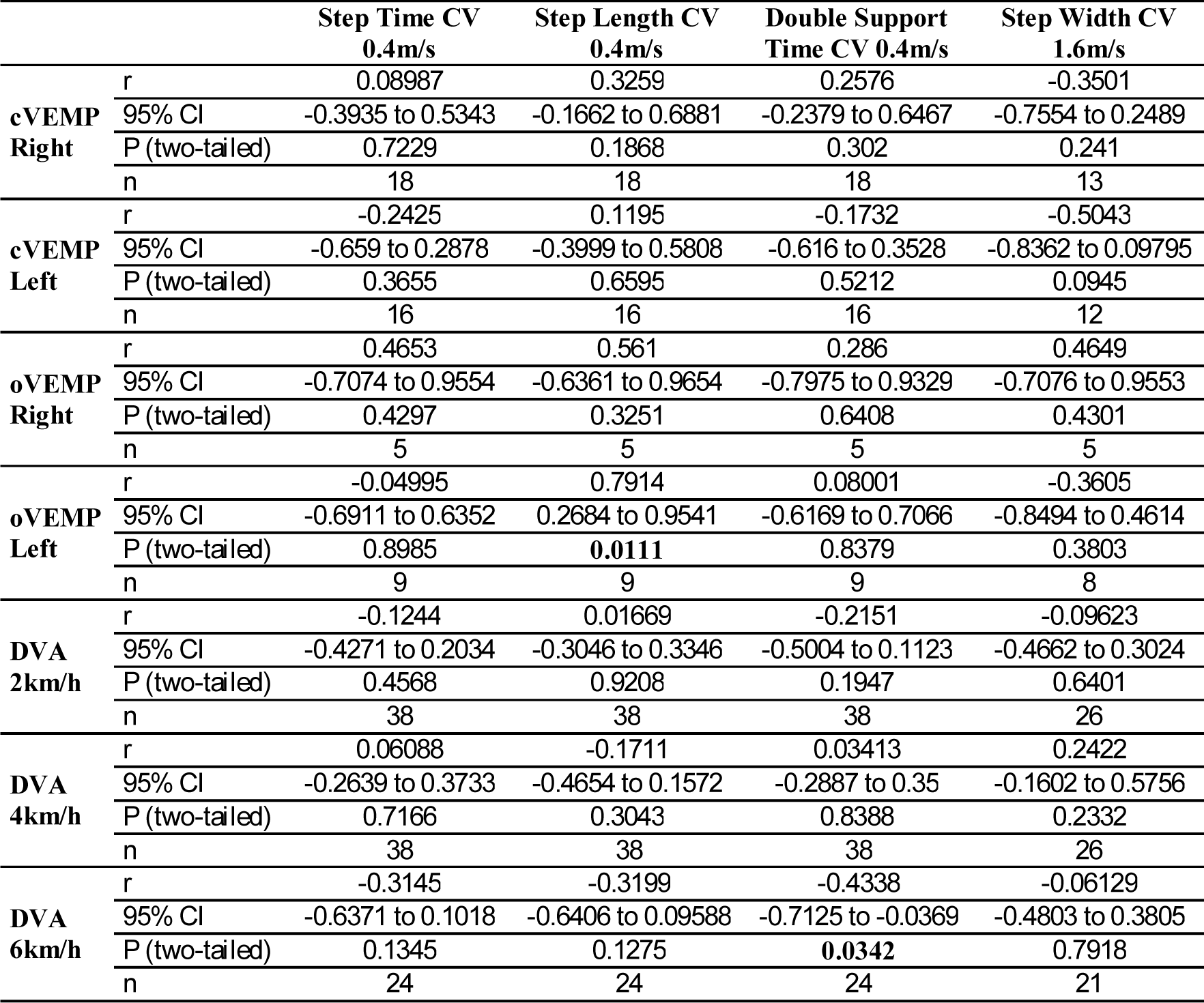
Pearson correlations between the cVEMP and oVEMP thresholds, the change in logMAR scores during each of the three DVA walking speeds and the gait parameters.

|  |  | **Step Time CV**<br>**0.4m/s** | **Step Length CV**<br>**0.4m/s** | **Double Support**<br>**Time CV 0.4m/s** | **Step Width CV**<br>**1.6m/s** |
| --- | --- | --- | --- | --- | --- |
| **cVEMP**<br>**Right** | r | 0.08987 | 0.3259 | 0.2576 | -0.3501 |
|  | 95% CI | -0.3935 to 0.5343 | -0.1662 to 0.6881 | -0.2379 to 0.6467 | -0.7554 to 0.2489 |
|  | P (two-tailed) | 0.7229 | 0.1868 | 0.302 | 0.241 |
|  | n | 18 | 18 | 18 | 13 |
| **cVEMP**<br>**Left** | r | -0.2425 | 0.1195 | -0.1732 | -0.5043 |
|  | 95% CI | -0.659 to 0.2878 | -0.3999 to 0.5808 | -0.616 to 0.3528 | -0.8362 to 0.09795 |
|  | P (two-tailed) | 0.3655 | 0.6595 | 0.5212 | 0.0945 |
|  | n | 16 | 16 | 16 | 12 |
| **oVEMP**<br>**Right** | r | 0.4653 | 0.561 | 0.286 | 0.4649 |
|  | 95% CI | -0.7074 to 0.9554 | -0.6361 to 0.9654 | -0.7975 to 0.9329 | -0.7076 to 0.9553 |
|  | P (two-tailed) | 0.4297 | 0.3251 | 0.6408 | 0.4301 |
|  | n | 5 | 5 | 5 | 5 |
| **oVEMP**<br>**Left** | r | -0.04995 | 0.7914 | 0.08001 | -0.3605 |
|  | 95% CI | -0.6911 to 0.6352 | 0.2684 to 0.9541 | -0.6169 to 0.7066 | -0.8494 to 0.4614 |
|  | P (two-tailed) | 0.8985 | **0.0111** | 0.8379 | 0.3803 |
|  | n | 9 | 9 | 9 | 8 |
| **DVA**<br>**2km/h** | r | -0.1244 | 0.01669 | -0.2151 | -0.09623 |
|  | 95% CI | -0.4271 to 0.2034 | -0.3046 to 0.3346 | -0.5004 to 0.1123 | -0.4662 to 0.3024 |
|  | P (two-tailed) | 0.4568 | 0.9208 | 0.1947 | 0.6401 |
|  | n | 38 | 38 | 38 | 26 |
| **DVA**<br>**4km/h** | r | 0.06088 | -0.1711 | 0.03413 | 0.2422 |
|  | 95% CI | -0.2639 to 0.3733 | -0.4654 to 0.1572 | -0.2887 to 0.35 | -0.1602 to 0.5756 |
|  | P (two-tailed) | 0.7166 | 0.3043 | 0.8388 | 0.2332 |
|  | n | 38 | 38 | 38 | 26 |
| **DVA**<br>**6km/h** | r | -0.3145 | -0.3199 | -0.4338 | -0.06129 |
|  | 95% CI | -0.6371 to 0.1018 | -0.6406 to 0.09588 | -0.7125 to -0.0369 | -0.4803 to 0.3805 |
|  | P (two-tailed) | 0.1345 | 0.1275 | **0.0342** | 0.7918 |
|  | n | 24 | 24 | 24 | 21 |

### Post-hoc Analysis of Gait Data based on VEMP Results

In order to further investigate differences within the patient group, we conducted an analysis of the gait data of the participants with and without at least one detected VEMP threshold for the same four parameters as the correlations: the CVs of step time, step length and double support time at 0.4m/s and the CV of step width at 1.6m/s. Given that all of the participants with no VEMP threshold detected also had abnormal outcomes on the vHIT for most or all of the six directions tested, the purpose of this analysis was to compare the gait of participants with and without detectable canal and otolith function. Independent samples t-tests with Welch’s corrections did not reveal any significant differences between the participants with and without at least one detectable VEMP threshold (0.0965<P<0.746).

## Discussion

In this study, we aimed to determine the effects of systematic increases in walking speed on spatiotemporal gait parameters and their variability in people with BVP. Specifically, we investigated if these parameters would distinguish between healthy participants and participants with BVP, and between patients with BVP who could and could not complete all of the planned walking speed trials (a simple proxy of locomotor capacity). Our hypothesis, that step and double support time and step length variability would systematically reduce with increases in walking speed, whereas step width variability would systematically increase, was confirmed as significant effects of walking speed were found for all gait variability parameters. We additionally hypothesized that step and double support time and step length variability at slower walking speeds would be most distinguishing between the healthy control participants and patients with BVP, and also between the patients with BVP that could complete the measurement protocol, and the patients with BVP that could only partially complete the measurement protocol, whereas step width variability would be most distinguishing between these groups at faster walking speeds. This hypothesis was partly confirmed; step length CV differed between groups BVP and Young and between groups BVP All Gait and BVP Part Gait, double support time CV differed between groups BVP and Young and step width CV differed between groups BVP and Young and BVP and Older for step width variability, but other parameters did not significantly differ at the pairwise comparison level, despite the group effects found for all parameters except step width CV in the BVP All Gait vs. BVP Part Gait analysis.

Our secondary aim was to conduct an explorative analysis in the patient groups by examining correlations between the outcomes of four clinical vestibular tests (vHIT, oVEMP, cVEMP, DVA) and the most distinguishing gait parameters identified. Only one significant correlations between the change in logMAR scores during the DVA and the gait parameters were found (6km/h and Double Support CV; Table 2). One significant correlation of 16 was found between the VEMP thresholds and the gait parameters, but only nine pairs of data were included in this test and if a Bonferroni correction is made for the p values of these 16 tests, it is no longer significant (oVEMP Left and Step Length CV at 0.4m/s; Table 2). Similarly, the one significant correlation between a DVA parameter and gait variability (DVA 6km/h and Double support time CV 0.4m/s) does not meet the significance threshold if a Bonferroni correction for the 12 tests is made. Even though this study clearly demonstrates the significant contribution of vestibular function to gait, our exploratory analysis confirms the complex contribution of vestibular information during every-day activities and the difficulty in translating current objective clinical measures to highly relevant patient symptoms.

Determining meaningful and distinguishing gait parameters in BVP is vital for the development of interventions, as is using tasks that sufficiently replicate the day-to-day challenges of these patients, in order to determine candidates for intervention and to assess the effect of those interventions. Two promising interventions currently under development and investigation include noisy galvanic vestibular stimulation (nGVS) and vestibular implants (Guinand et al. 2015; Guyot et al. 2016; Lewis 2016; Perez Fornos et al. 2017; Wuehr et al. 2017). Discussions of these treatment options can be found elsewhere (Guyot et al. 2016; Wuehr et al. 2017), but in the context of this study, it is important to note that both options show early signs of utility for improving the gait of people with BVP (McCrum et al. 2016; Wuehr et al. 2016). However, it remains to be seen if improvement due to nGVS or a vestibular implant in steady state gait would likewise be seen in more dynamic locomotor task performance, where even unilateral vestibulopathy leads to significantly poorer stability performance (McCrum et al. 2014). Related to this, it should be noted that while this study examined spatiotemporal variability, differences in dynamic gait stability were not directly assessed and the two are not necessarily equivalent (Bruijn et al. 2013; Dingwell et al. 2001; Perry and Srinivasan 2017). The parameters presented here represent the amount of variability in particular gait parameters, but do not necessarily indicate the overall stability of the participants. Therefore, future work should investigate how dynamic gait stability is altered in BVP and how this is affected by changes in walking speed.

The current study confirmed previous findings of reductions in temporal gait variability and reductions in sagittal plane spatial gait variability in vestibulopathy during faster, compared to slower walking (Schniepp et al. 2017; Schniepp et al. 2012; Wuehr et al. 2016). We extend these previous findings as the current study employed fixed (not self-selected) speeds that were systematically increased, with 120 steps analyzed per speed, thereby improving the reliability of the outcomes. Importantly, the current results further the previous findings by additionally showing that these parameters are related to the locomotor capacities of people with BVP.

We also confirmed previously reported increases in step width variability with increasing walking speeds in people with BVP (Wuehr et al. 2016). Previous studies have shown that vestibular perturbations have less impact on direction and variability at higher walking speeds (Fitzpatrick et al. 1999; Jahn et al. 2000) and that the vestibular influence on lower limb muscles is selectively suppressed with increased cadence and speed during walking (Dakin et al. 2013; Forbes et al. 2017). However, the current step width variability results, combined with those of (Wuehr et al. 2016) suggest that vestibular information remains important for mediolateral foot placement at increased walking speeds. During the swing phase when foot placement is coordinated and determined, there is reduced proprioceptive input due to only one foot being in contact with the ground. Therefore, we could reason that vestibular input becomes more important in this phase, and disturbed or lacking vestibular input may therefore decrease the accuracy of foot placement. These results also provide some explanation as to why people with BVP do not self-select walking speeds that minimize temporal or sagittal plane spatial gait variability (Schniepp et al. 2017; Schniepp et al. 2012; Wuehr et al. 2016). Dramatic increases in step width variability may be undesirable due to reduced stability control or increased energetic costs of mediolateral stabilization (Dean et al. 2007; Donelan et al. 2004; O’Connor et al. 2012). Based on the current results, either reason is plausible, as some of the participants in the BVP Part Gait group did not continue to the faster speeds due to instability, while others could not continue due to being unable to keep up with the speed of the treadmill (implying an energetic or physiological limitation, not a stability-related one). The vestibular influence on gait economy has not yet, to our knowledge, been thoroughly investigated, and is therefore an area for future research.

The healthy control groups in this study were not directly age matched with the BVP group, but rather represent healthy participants at the younger and older end of the age range of the BVP group. In the current results, the variability in step time, double support time and step length of the older group tends to fall between that of the younger and BVP group, showing few statistical differences to either, although we suspect that this is due to a lack of statistical power at the pairwise comparison level. The boxplots seem to indicate that the group Older tend towards the results of group Young for double support time and step length variability. However, the group difference in step width variability appear to be more robust, with large significant differences between the BVP group and each healthy group, and no difference due to healthy ageing age alone, in agreement with previous studies (Herssens et al. 2018; Hollman et al. 2011). However, other limitations in the current study should be kept in mind. Caution should be taken in comparing the CV of step width to studies of overground walking, as it has been shown previously in healthy participants that walking on the CAREN results in increased step width variability compared to overground walking (Gates et al. 2012). Additionally, treadmill walking appears to be more challenging than overground walking for people with BVP, evidenced by the fact that the BVP No Gait group were not able to successfully complete the familiarization period, despite reporting being able to walking independently without assistance. We would therefore caution a direct comparison of treadmill-derived gait results with overground gait results in BVP. Regarding the fact that the healthy groups walked with optic flow and the BVP group walked with the virtual environment fixed (so as to provide the same lighting), we do not expect that this difference would have altered our results, as two previous studies found no, or negligible, differences in the parameters assessed here between fixed speed walking with and without virtual reality (Katsavelis et al. 2010; Sloot et al. 2014). The only previous study that did find differences in gait variability due to virtual reality that we are aware of is that of Hollman et al. (2006). However, Hollman et al. (2006) used an insufficient number of data points to reliably assess gait variability (Katsavelis et al. 2010) and used a substantially different virtual reality setup to the current study. We used a setup comparable to that of Sloot et al. (2014), who found no differences in gait variability as a result of using a virtual reality screen with optic flow. Finally, the effect sizes of the difference in step width variability with and without virtual reality and optic flow from Hollman et al. (2006) are much smaller than those found in the current study between Young and BVP All Gait groups at similar walking speeds (Cohen’s d of 0.238-0.657 in Hollman et al. (2006) vs. 1.064-1.382 in the current study).

We also acknowledge that our division of participants into the BVP All Gait and BVP Part Gait groups is based on a rather simple criterion. Of the 12 participants in the BVP Part Gait group, one participant stopped after 0.8m/s, one after 1.0m/s, four after 1.2m/s and six after 1.4m/s and therefore, the range of locomotor capacities within this group is likely broad. Reasons for lack of completion also varied across the participants, with some stopping due to lack of stability control (too much lateral deviation with a risk of stepping off the treadmill) and others unable to keep up with a faster belt speed. Nevertheless, we found significant group effects on gait variability, indicating the potential association between gait variability and overall locomotor capacity in BVP. Further research into gait parameters that can distinguish between patients with different functional limitations is encouraged to aid the development of accurate diagnostic functional testing protocols.

In conclusion, spatiotemporal gait parameters and their variability show speed-dependency in people with BVP and in healthy adults. In particular, step length variability at slower speeds and step width variability at faster speeds were the most distinguishing parameters between the healthy participants and people with BVP, and within people with BVP who have different locomotor capacities. Gait variability in BVP was generally not correlated with the clinical tests of vestibular function. The current findings indicate that analysis of gait variability at multiple speeds has potential as an assessment tool for vestibular interventions.

## Acknowledgements

The authors thank Wouter Bijnens, Rachel Senden and Rik Marcellis for their support with the measurements.

## Author Contributions

Conception of the study: CM, RvdB, KK, HK, KM. Data Collection: CM, FL. Data Analysis: CM, PW, FL. Interpreted results: All authors. Prepared Figures: CM. Drafted the article: CM. Reviewed and revised the article: All authors. Approved final version: All authors.

## References

Bailey IL, and Lovie JE. New design principles for visual acuity letter charts. Am J Optom Physiol Opt 53: 740–745, 1976.

Bauby CE, and Kuo AD. Active control of lateral balance in human walking. J Biomech 33: 1433–1440, 2000.

Brandt T, Strupp M, and Benson J. You are better off running than walking with acute vestibulopathy. Lancet 354: 746, 1999.

Brantberg K, and Lofqvist L. Preserved vestibular evoked myogenic potentials (VEMP) in some patients with walking-induced oscillopsia due to bilateral vestibulopathy. J Vestib Res 17: 33–38, 2007.

Bruijn SM, Meijer OG, Beek PJ, and van Dieen JH. Assessing the stability of human locomotion: a review of current measures. Journal of the Royal Society, Interface / the Royal Society 10: 20120999, 2013.

Dakin CJ, Inglis JT, Chua R, and Blouin JS. Muscle-specific modulation of vestibular reflexes with increased locomotor velocity and cadence. J Neurophysiol 110: 86–94, 2013.

Dean JC, Alexander NB, and Kuo AD. The effect of lateral stabilization on walking in young and old adults. IEEE Trans Biomed Eng 54: 1919–1926, 2007.

Dingwell JB, Cusumano JP, Cavanagh PR, and Sternad D. Local dynamic stability versus kinematic variability of continuous overground and treadmill walking. J Biomech Eng 123: 27–32, 2001.

Donelan JM, Shipman DW, Kram R, and Kuo AD. Mechanical and metabolic requirements for active lateral stabilization in human walking. J Biomech 37: 827–835, 2004.

Fitzpatrick RC, Wardman DL, and Taylor JL. Effects of galvanic vestibular stimulation during human walking. J Physiol 517 (Pt 3): 931–939, 1999.

Forbes PA, Vlutters M, Dakin CJ, van der Kooij H, Blouin JS, and Schouten AC. Rapid limb-specific modulation of vestibular contributions to ankle muscle activity during locomotion. J Physiol 595: 2175–2195, 2017.

Fujimoto C, Murofushi T, Chihara Y, Suzuki M, Yamasoba T, and Iwasaki S. Novel subtype of idiopathic bilateral vestibulopathy: bilateral absence of vestibular evoked myogenic potentials in the presence of normal caloric responses. J Neurol 256: 1488–1492, 2009.

Gates DH, Darter BJ, Dingwell JB, and Wilken JM. Comparison of walking overground and in a Computer Assisted Rehabilitation Environment (CAREN) in individuals with and without transtibial amputation. J Neuroeng Rehabil 9: 2012.

Govender S, Rosengren SM, and Colebatch JG. Vestibular neuritis has selective effects on air- and bone-conducted cervical and ocular vestibular evoked myogenic potentials. Clin Neurophysiol 122: 1246–1255, 2011.

Guinand N, Boselie F, Guyot JP, and Kingma H. Quality of life of patients with bilateral vestibulopathy. Ann Otol Rhinol Laryngol 121: 471–477, 2012a.

Guinand N, Pijnenburg M, Janssen M, and Kingma H. Visual Acuity While Walking and Oscillopsia Severity in Healthy Subjects and Patients With Unilateral and Bilateral Vestibular Function Loss. Archives of Otolaryngology-Head & Neck Surgery 138: 301–306, 2012b.

Guinand N, van de Berg R, Cavuscens S, Stokroos RJ, Ranieri M, Pelizzone M, Kingma H, Guyot JP, and Perez-Fornos A. Vestibular Implants: 8 Years of Experience with Electrical Stimulation of the Vestibular Nerve in 11 Patients with Bilateral Vestibular Loss. ORL J Otorhinolaryngol Relat Spec 77: 227–240, 2015.

Guyot JP, Perez Fornos A, Guinand N, van de Berg R, Stokroos R, and Kingma H. Vestibular assistance systems: promises and challenges. J Neurol 263 Suppl 1: S30–35, 2016.

Herssens N, Verbecque E, Hallemans A, Vereeck L, Van Rompaey V, and Saeys W. Do spatiotemporal parameters and gait variability differ across the lifespan of healthy adults? A systematic review. Gait Posture 64: 181–190, 2018.

Hollman JH, Brey RH, Robb RA, Bang TJ, and Kaufman KR. Spatiotemporal gait deviations in a virtual reality environment. Gait Posture 23: 441–444, 2006.

Hollman JH, McDade EM, and Petersen RC. Normative spatiotemporal gait parameters in older adults. Gait Posture 34: 111–118, 2011.

Horak FB, Kluzik J, and Hlavacka F. Velocity dependence of vestibular information for postural control on tilting surfaces. J Neurophysiol 116: 1468–1479, 2016.

Jahn K, Strupp M, Schneider E, Dieterich M, and Brandt T. Differential effects of vestibular stimulation on walking and running. Neuroreport 11: 1745–1748, 2000.

Jahn K, Strupp M, Schneider E, Dieterich M, and Brandt T. Visually induced gait deviations during different locomotion speeds. Exp Brain Res 141: 370–374, 2001.

Katsavelis D, Mukherjee M, Decker L, and Stergiou N. The effect of virtual reality on gait variability. Nonlinear Dynamics Psychol Life Sci 14: 239–256, 2010.

Lewis RF. Vestibular implants studied in animal models: clinical and scientific implications. J Neurophysiol 116: 2777–2788, 2016.

Lucieer F, Vonk P, Guinand N, Stokroos R, Kingma H, and van de Berg R. Bilateral Vestibular Hypofunction: Insights in Etiologies, Clinical Subtypes, and Diagnostics. Front Neurol 7: 26, 2016.

McCrum C, Eysel-Gosepath K, Epro G, Meijer K, Savelberg HH, Brüggemann GP, and Karamanidis K. Deficient recovery response and adaptive feedback potential in dynamic gait stability in unilateral peripheral vestibular disorder patients. Physiol Rep 2: e12222, 2014.

McCrum C, Willems P, van de Berg R, Cavuscens S, Guinand N, Guyot JP, Perez Fornos A, Marcellis R, Ranieri M, Senden R, Stokroos R, Zijlstra W, Karamanidis K, Meijer K, and Kingma H. Preliminary observations of the acute effects of vestibular nerve stimulation on stride length and time in two patients with bilateral vestibular hypofunction. Gait Posture 49: 124, 2016.

McGarvie LA, MacDougall HG, Halmagyi GM, Burgess AM, Weber KP, and Curthoys IS. The Video Head Impulse Test (vHIT) of Semicircular Canal Function - Age-Dependent Normative Values of VOR Gain in Healthy Subjects. Front Neurol 6: 154, 2015.

O’Connor SM, Xu HZ, and Kuo AD. Energetic cost of walking with increased step variability. Gait Posture 36: 102–107, 2012.

Perez Fornos A, Cavuscens S, Ranieri M, van de Berg R, Stokroos R, Kingma H, Guyot JP, and Guinand N. The vestibular implant: A probe in orbit around the human balance system. J Vestib Res 27: 51–61, 2017.

Perry JA, and Srinivasan M. Walking with wider steps changes foot placement control, increases kinematic variability and does not improve linear stability. Royal Society open science 4: 160627, 2017.

Piker EG, Jacobson GP, Burkard RF, McCaslin DL, and Hood LJ. Effects of age on the tuning of the cVEMP and oVEMP. Ear Hear 34: e65–73, 2013.

Schlick C, Schniepp R, Loidl V, Wuehr M, Hesselbarth K, and Jahn K. Falls and fear of falling in vertigo and balance disorders: A controlled cross-sectional study. J Vestib Res 25: 241–251, 2016.

Schniepp R, Schlick C, Schenkel F, Pradhan C, Jahn K, Brandt T, and Wuehr M. Clinical and neurophysiological risk factors for falls in patients with bilateral vestibulopathy. J Neurol 264: 277– 283, 2017.

Schniepp R, Wuehr M, Neuhaeusser M, Kamenova M, Dimitriadis K, Klopstock T, Strupp M, Brandt T, and Jahn K. Locomotion speed determines gait variability in cerebellar ataxia and vestibular failure. Mov Disord 27: 125–131, 2012.

Sloan LL. New test charts for the measurement of visual acuity at far and near distances. Am J Ophthalmol 48: 807–813, 1959.

Sloot LH, van der Krogt MM, and Harlaar J. Effects of adding a virtual reality environment to different modes of treadmill walking. Gait Posture 39: 939–945, 2014.

Sprenger A, Wojak JF, Jandl NM, and Helmchen C. Postural Control in Bilateral Vestibular Failure: Its Relation to Visual, Proprioceptive, Vestibular, and Cognitive Input. Front Neurol 8: 444, 2017.

Strupp M, Kim JS, Murofushi T, Straumann D, Jen JC, Rosengren SM, Della Santina CC, and Kingma H. Bilateral vestibulopathy: Diagnostic criteria Consensus document of the Classification Committee of the Barany Society. J Vestib Res 27: 177–189, 2017.

Valko Y, Rosengren SM, Jung HH, Straumann D, Landau K, and Weber KP. Ocular vestibular evoked myogenic potentials as a test for myasthenia gravis. Neurology 86: 660–668, 2016.

van de Berg R, van Tilburg M, and Kingma H. Bilateral Vestibular Hypofunction: Challenges in Establishing the Diagnosis in Adults. ORL J Otorhinolaryngol Relat Spec 77: 197–218, 2015.

van Dooren TS, Lucieer FMP, Janssen AML, Kingma H, and van de Berg R. The Video Head Impulse Test and the Influence of Daily Use of Spectacles to Correct a Refractive Error. Front Neurol 9: 125, 2018.

van Tilburg MJ, Herrmann BS, Guinan JJ, Jr., and Rauch SD. Increasing the Stimulation Rate Reduces cVEMP Testing Time by More Than Half With No Significant Difference in Threshold. Otol Neurotol 37: 933–936, 2016.

Wuehr M, Decker J, and Schniepp R. Noisy galvanic vestibular stimulation: an emerging treatment option for bilateral vestibulopathy. J Neurol 264: 81–86, 2017.

Wuehr M, Nusser E, Decker J, Krafczyk S, Straube A, Brandt T, Jahn K, and Schniepp R. Noisy vestibular stimulation improves dynamic walking stability in bilateral vestibulopathy. Neurology 2016.

Wuehr M, Schniepp R, Pradhan C, Ilmberger J, Strupp M, Brandt T, and Jahn K. Differential effects of absent visual feedback control on gait variability during different locomotion speeds. Exp Brain Res 224: 287–294, 2013.

Yip CW, Glaser M, Frenzel C, Bayer O, and Strupp M. Comparison of the Bedside Head-Impulse Test with the Video Head-Impulse Test in a Clinical Practice Setting: A Prospective Study of 500 Outpatients. Front Neurol 7: 58, 2016.

Zeni JA, Jr., Richards JG, and Higginson JS. Two simple methods for determining gait events during treadmill and overground walking using kinematic data. Gait Posture 27: 710–714, 2008.

