## Supplementary for "Is faster always better? The walking speed-dependency of gait variability in bilateral vestibulopathy"

**Supplementary Table 1: Bonferroni pairwise between group comparison results for the means of the gait parameters**

| Walking Speed (m/s) | Groups | Step Time Means | | | Step Length Means | | | Double Support Time Means | | | Step Width Means | | |
|---|---|---|---|---|---|---|---|---|---|---|---|---|---|
| | | **Mean Diff.** | **95.00% CI of diff.** | **Adjusted P Value** | **Mean Diff.** | **95.00% CI of diff.** | **Adjusted P Value** | **Mean Diff.** | **95.00% CI of diff.** | **Adjusted P Value** | **Mean Diff.** | **95.00% CI of diff.** | **Adjusted P Value** |
| 0.4 | BVP vs. Young | -0.1579 | -0.2419 to -0.07384 | **0.0001** | -0.06307 | -0.09654 to -0.02960 | **0.0001** | -0.07791 | -0.1170 to -0.03885 | **<0.0001** | 0.03707 | 0.003385 to 0.07075 | **0.0268** |
| | BVP vs. Older | -0.09576 | -0.1938 to 0.002244 | 0.057 | -0.03835 | -0.07742 to 0.0007265 | 0.0556 | -0.02706 | -0.07522 to 0.02110 | 0.4786 | 0.03175 | -0.009976 to 0.07347 | 0.1867 |
| | Young vs. Older | 0.06212 | -0.04178 to 0.1660 | 0.4046 | 0.02472 | -0.01663 to 0.06607 | 0.4044 | 0.05086 | 0.0008404 to 0.1009 | **0.0454** | -0.00532 | -0.04651 to 0.03587 | >0.9999 |
| 0.6 | BVP vs. Young | -0.09504 | -0.1465 to -0.04361 | **0.0002** | -0.05714 | -0.08806 to -0.02622 | **0.0002** | -0.03592 | -0.05997 to -0.01188 | **0.0024** | 0.02883 | -0.006725 to 0.06439 | 0.1482 |
| | BVP vs. Older | -0.05925 | -0.1277 to 0.009149 | 0.1047 | -0.03606 | -0.07720 to 0.005079 | 0.099 | -0.01187 | -0.03891 to 0.01517 | 0.7921 | 0.03253 | -0.01095 to 0.07602 | 0.2005 |
| | Young vs. Older | 0.03579 | -0.03479 to 0.1064 | 0.5979 | 0.02108 | -0.02143 to 0.06358 | 0.627 | 0.02405 | -0.006358 to 0.05446 | 0.1572 | 0.003702 | -0.03852 to 0.04592 | >0.9999 |
| 0.8 | BVP vs. Young | -0.05854 | -0.09343 to -0.02365 | **0.0004** | -0.04711 | -0.07525 to -0.01898 | **0.0005** | -0.01993 | -0.03893 to -0.0009376 | **0.0374** | 0.01668 | -0.02021 to 0.05358 | 0.7954 |
| | BVP vs. Older | -0.0398 | -0.08653 to 0.006931 | 0.1149 | -0.03217 | -0.06972 to 0.005381 | 0.1118 | -0.009608 | -0.02740 to 0.008184 | 0.5388 | 0.02243 | -0.01548 to 0.06034 | 0.4394 |
| | Young vs. Older | 0.01874 | -0.02655 to 0.06403 | 0.8688 | 0.01494 | -0.02150 to 0.05138 | 0.8821 | 0.01033 | -0.01040 to 0.03105 | 0.6294 | 0.005752 | -0.03036 to 0.04187 | >0.9999 |
| 1 | BVP vs. Young | -0.05273 | -0.08212 to -0.02334 | **0.0002** | -0.05325 | -0.08276 to -0.02374 | **0.0002** | -0.0124 | -0.02665 to 0.001852 | 0.1062 | 0.01449 | -0.02451 to 0.05349 | >0.9999 |
| | BVP vs. Older | -0.02875 | -0.06112 to 0.003616 | 0.0959 | -0.02911 | -0.06122 to 0.002991 | 0.0864 | -0.00682 | -0.02207 to 0.008430 | 0.7972 | 0.0166 | -0.02325 to 0.05645 | 0.9078 |
| | Young vs. Older | 0.02398 | -0.006335 to 0.05429 | 0.1569 | 0.02413 | -0.006074 to 0.05434 | 0.1507 | 0.005578 | -0.009967 to 0.02112 | >0.9999 | 0.002111 | -0.03669 to 0.04091 | >0.9999 |
| 1.2 | BVP vs. Young | -0.03527 | -0.05426 to -0.01627 | **0.0002** | -0.04224 | -0.06470 to -0.01978 | **0.0001** | -0.004739 | -0.01834 to 0.008861 | >0.9999 | 0.01157 | -0.02591 to 0.04906 | >0.9999 |
| | BVP vs. Older | -0.01281 | -0.03491 to 0.009291 | 0.4441 | -0.01505 | -0.04133 to 0.01123 | 0.4584 | -0.001326 | -0.01452 to 0.01187 | >0.9999 | 0.01897 | -0.02107 to 0.05900 | 0.7251 |
| | Young vs. Older | 0.02246 | -0.0008739 to 0.04579 | 0.0618 | 0.02719 | -0.0004576 to 0.05483 | 0.0549 | 0.003413 | -0.01246 to 0.01929 | >0.9999 | 0.007391 | -0.03035 to 0.04514 | >0.9999 |
| 1.4 | BVP vs. Young | -0.02955 | -0.04798 to -0.01112 | **0.001** | -0.04111 | -0.06684 to -0.01537 | **0.001** | -0.002945 | -0.01446 to 0.008574 | >0.9999 | 0.01293 | -0.02312 to 0.04897 | >0.9999 |
| | BVP vs. Older | -0.009134 | -0.02823 to 0.009961 | 0.7002 | -0.01253 | -0.03915 to 0.01409 | 0.7227 | -0.003735 | -0.01521 to 0.007745 | >0.9999 | 0.009436 | -0.03317 to 0.05204 | >0.9999 |
| | Young vs. Older | 0.02042 | 0.0001919 to 0.04065 | **0.0474** | 0.02858 | 0.0003932 to 0.05676 | **0.0461** | -0.00079 | -0.01374 to 0.01216 | >0.9999 | -0.00349 | -0.04146 to 0.03448 | >0.9999 |
| 1.6 | BVP vs. Young | -0.01865 | -0.03602 to -0.001278 | **0.0324** | -0.02908 | -0.05752 to -0.0006455 | **0.0438** | -0.001623 | -0.01276 to 0.009513 | >0.9999 | 0.005952 | -0.03517 to 0.04707 | >0.9999 |
| | BVP vs. Older | -0.006196 | -0.02836 to 0.01597 | >0.9999 | -0.01025 | -0.04572 to 0.02522 | >0.9999 | -0.002359 | -0.01579 to 0.01108 | >0.9999 | 0.002257 | -0.03830 to 0.04282 | >0.9999 |
| | Young vs. Older | 0.01245 | -0.01100 to 0.03591 | 0.5373 | 0.01883 | -0.01903 to 0.05669 | 0.6219 | -0.000736 | -0.01479 to 0.01332 | >0.9999 | -0.003695 | -0.03929 to 0.03190 | >0.9999 |

**Supplementary Table 2: Bonferroni pairwise between group comparison results for the coefficients of variation of the gait parameters**

| Walking Speed (m/s) | Groups | Step Time CV | | | Step Length CV | | | Double Support Time CV | | | Step Width CV | | |
|---|---|---|---|---|---|---|---|---|---|---|---|---|---|
| | | Mean Diff. | 95.00% CI of diff. | Adjusted P Value | Mean Diff. | 95.00% CI of diff. | Adjusted P Value | Mean Diff. | 95.00% CI of diff. | Adjusted P Value | Mean Diff. | 95.00% CI of diff. | Adjusted P Value |
| 0.4 | BVP vs. Young | 1.271 | -0.09019 to 2.631 | 0.073 | 2.791 | 0.8092 to 4.773 | **0.004** | 4.523 | 1.343 to 7.704 | **0.0033** | 0.7604 | -2.313 to 3.834 | >0.9999 |
| | BVP vs. Older | 0.3193 | -1.749 to 2.388 | >0.9999 | 1.611 | -1.029 to 4.252 | 0.3731 | 1.269 | -4.099 to 6.637 | >0.9999 | 1.46 | -3.380 to 6.299 | >0.9999 |
| | Young vs. Older | -0.9513 | -3.125 to 1.222 | 0.79 | -1.18 | -3.990 to 1.630 | 0.8556 | -3.254 | -8.710 to 2.201 | 0.3927 | 0.6991 | -4.033 to 5.432 | >0.9999 |
| 0.6 | BVP vs. Young | 1.355 | 0.4934 to 2.216 | **0.0009** | 2.699 | 1.153 to 4.246 | **0.0004** | 3.953 | 1.783 to 6.123 | **0.0001** | 4.984 | 0.9730 to 8.994 | **0.0102** |
| | BVP vs. Older | 0.9247 | -0.5926 to 2.442 | 0.3752 | 1.704 | -0.3164 to 3.724 | 0.1182 | 2.94 | -1.040 to 6.920 | 0.1989 | 4.169 | -1.194 to 9.531 | 0.1745 |
| | Young vs. Older | -0.4301 | -1.909 to 1.049 | >0.9999 | -0.9957 | -3.101 to 1.109 | 0.691 | -1.013 | -4.941 to 2.915 | >0.9999 | -0.8149 | -5.417 to 3.787 | >0.9999 |
| 0.8 | BVP vs. Young | 1.201 | 0.5513 to 1.850 | **<0.0001** | 2.606 | 1.459 to 3.754 | **<0.0001** | 3.473 | 1.361 to 5.586 | **0.0006** | 10.67 | 4.704 to 16.64 | **0.0002** |
| | BVP vs. Older | 0.5263 | -0.6012 to 1.654 | 0.7134 | 1.15 | -0.3300 to 2.630 | 0.1724 | 2.72 | 0.03094 to 5.410 | **0.0468** | 6.319 | -3.386 to 16.02 | 0.3216 |
| | Young vs. Older | -0.6742 | -1.725 to 0.3768 | 0.3065 | -1.456 | -2.859 to -0.05344 | **0.0404** | -0.7528 | -3.407 to 1.901 | >0.9999 | -4.355 | -12.93 to 4.216 | 0.5519 |
| 1 | BVP vs. Young | 0.8551 | 0.01044 to 1.700 | **0.0466** | 2.091 | 0.9474 to 3.234 | **0.0001** | 3.203 | 0.5901 to 5.815 | **0.012** | 12.86 | 6.892 to 18.83 | **<0.0001** |
| | BVP vs. Older | 0.2881 | -0.5151 to 1.091 | >0.9999 | 0.8791 | -0.4269 to 2.185 | 0.2971 | 1.236 | -1.258 to 3.731 | 0.6655 | 11.18 | 4.064 to 18.29 | **0.001** |
| | Young vs. Older | -0.567 | -1.513 to 0.3790 | 0.4038 | -1.212 | -2.416 to -0.007390 | **0.0483** | -1.966 | -4.525 to 0.5927 | 0.177 | -1.681 | -6.651 to 3.289 | >0.9999 |
| 1.2 | BVP vs. Young | 0.6428 | 0.1572 to 1.128 | **0.0058** | 1.579 | 0.7094 to 2.448 | **0.0002** | 2.907 | 0.8770 to 4.937 | **0.0027** | 14.37 | 8.318 to 20.42 | **<0.0001** |
| | BVP vs. Older | 0.2557 | -0.3420 to 0.8533 | 0.8616 | 0.7506 | -0.01438 to 1.515 | 0.056 | 1.651 | -0.1934 to 3.495 | 0.0932 | 11.22 | 3.168 to 19.27 | **0.0038** |
| | Young vs. Older | -0.3872 | -0.9284 to 0.1541 | 0.2271 | -0.8281 | -1.647 to -0.009037 | **0.047** | -1.256 | -2.748 to 0.2368 | 0.1194 | -3.15 | -9.550 to 3.250 | 0.6027 |
| 1.4 | BVP vs. Young | 0.7897 | 0.3740 to 1.205 | **<0.0001** | 1.492 | 0.7862 to 2.198 | **<0.0001** | 2.492 | 0.1591 to 4.824 | **0.0329** | 15.22 | 7.806 to 22.63 | **<0.0001** |
| | BVP vs. Older | 0.4487 | -0.02622 to 0.9237 | 0.0695 | 0.6927 | 0.01951 to 1.366 | **0.0419** | 1.558 | -0.6551 to 3.771 | 0.2584 | 13.38 | 4.020 to 22.74 | **0.0029** |
| | Young vs. Older | -0.341 | -0.7297 to 0.04760 | 0.0983 | -0.7993 | -1.422 to -0.1769 | **0.0092** | -0.9334 | -3.036 to 1.170 | 0.7864 | -1.842 | -9.031 to 5.347 | >0.9999 |
| 1.6 | BVP vs. Young | 0.4406 | 0.06548 to 0.8158 | **0.0169** | 0.923 | 0.2817 to 1.564 | **0.0029** | 2.356 | 0.3356 to 4.376 | **0.0178** | 13.11 | 5.274 to 20.95 | **0.0005** |
| | BVP vs. Older | 0.1737 | -0.2706 to 0.6179 | 0.9734 | 0.3466 | -0.2665 to 0.9596 | 0.4899 | 1.059 | -1.125 to 3.243 | 0.6811 | 14.73 | 7.518 to 21.94 | **<0.0001** |
| | Young vs. Older | -0.267 | -0.7029 to 0.1690 | 0.3732 | -0.5765 | -1.174 to 0.02123 | 0.0611 | -1.297 | -3.418 to 0.8240 | 0.3782 | 1.619 | -3.208 to 6.447 | >0.9999 |

### Supplementary Table 3: Bonferroni pairwise between BVP All Gait vs. BVP Part Gait comparison results for the coefficients of variation of the gait parameters

| Walking Speed (m/s) | Step Time CV | | | Step Length CV | | | Double Support Time CV | | | Step Width CV | | |
|---|---|---|---|---|---|---|---|---|---|---|---|---|
| | **Mean Diff.** | **95.00% CI of diff.** | **Adjusted P Value** | **Mean Diff.** | **95.00% CI of diff.** | **Adjusted P Value** | **Mean Diff.** | **95.00% CI of diff.** | **Adjusted P Value** | **Mean Diff.** | **95.00% CI of diff.** | **Adjusted P Value** |
| 0.4 | -1.572 | -3.820 to 0.6765 | 0.3059 | -3.163 | -6.032 to -0.2933 | **0.0259** | -4.748 | -11.28 to 1.787 | 0.2531 | -0.4195 | -6.080 to 5.241 | >0.9999 |
| 0.6 | -1.259 | -3.216 to 0.6981 | 0.4225 | -2.432 | -5.321 to 0.4568 | 0.1305 | -2.8 | -7.624 to 2.024 | 0.5944 | -1.423 | -11.73 to 8.881 | >0.9999 |
| 0.8 | -1.241 | -2.925 to 0.4423 | 0.2441 | -1.864 | -4.438 to 0.7090 | 0.2652 | -3.054 | -7.209 to 1.100 | 0.2551 | 1.35 | -12.73 to 15.42 | >0.9999 |
| 1 | -0.7285 | -2.039 to 0.5822 | 0.684 | -1.316 | -4.399 to 1.768 | >0.9999 | -3.068 | -9.497 to 3.361 | 0.9497 | -1.021 | -16.51 to 14.47 | >0.9999 |
| 1.2 | -0.3443 | -1.341 to 0.6527 | >0.9999 | -0.202 | -1.677 to 1.273 | >0.9999 | -1.29 | -5.509 to 2.930 | >0.9999 | -6.166 | -23.90 to 11.57 | >0.9999 |
| 1.4 | -1.089 | -2.639 to 0.4610 | 0.212 | -1.134 | -3.709 to 1.440 | 0.8482 | -5.092 | -15.66 to 5.471 | 0.6228 | -8.85 | -39.64 to 21.94 | >0.9999 |
